# Comparative Analysis of Global Hepatic Gene Expression in Adolescents and Adults with Non-alcoholic Fatty Liver Disease

**DOI:** 10.1101/2022.12.08.519606

**Authors:** Samer Gawrieh, Rebekah Karns, David E. Kleiner, Michael Olivier, Todd Jenkins, Thomas H. Inge, Naga P. Chalasani, Stavra Xanthakos

## Abstract

**Introduction:** To gain insights into the mechanisms underlying distinct nonalcoholic fatty liver disease (NAFLD) histological phenotypes between children and adults, we compared hepatic gene expression profiles associated with NAFLD phenotypes between the two age groups.

**Methods:** Histological characteristics of intra-operative liver biopsies from adolescents and adults undergoing bariatric surgery were assessed by the same pathologist using the non-alcoholic steatohepatitis (NASH) Clinical Research Network scoring system. Hepatic gene expression was measured by microarray analysis. Transcriptomic signatures of histological phenotypes between the two groups were compared, with significance defined as p-value <0.05 and a fold change >1.5.

**Results:** In 67 adolescents and 76 adults, distribution of histological phenotypes was: not-NAFLD (controls) 51% vs 39%, NAFL 39% vs 37%, and NASH 10% vs 24%, respectively. There were 279 differentially expressed genes in adolescents and 213 in adults with NAFLD vs controls. In adolescents, transcriptomes for NAFL vs controls, and borderline vs definite NASH were undifferentiable, whereas in adults, NAFL and borderline NASH demonstrated a transcriptomic gradient between controls and definite NASH. When applied to adolescents, significant adult genes discriminated borderline and definite NASH from control and NAFL, but the majority of significant pediatric genes were not portable to adults. Genes associated with NASH in adolescents and adults showed some ontological consistency but notable differences.

**Conclusions:** There is some similarity but major differences in the transcriptomic profiles associated with NAFLD between adolescents and adults with severe obesity. These data suggest different mechanisms contribute to the pathogenesis of NAFLD severity at different stages in life.

**Study Highlights:** *WHAT IS KNOWN:*

- Nonalcoholic fatty liver disease (NAFLD) is the most common liver disease in children and adults
- NAFLD histological features and severity differ between children and adults but reasons for these differences are not clear.

*WHAT IS NEW HERE:*

- Comparison of hepatic gene expression profiles between adolescents and adults with severe obesity and NAFLD showed some similarities but major differences in expressed genes
- The findings suggest the mechanisms driving different NAFLD severity and phenotypes are different at different stages in life.

## Introduction

Nonalcoholic fatty liver disease (NAFLD) is currently the leading global cause of chronic liver disease in children and adults^1-3^. NAFLD is estimated to affect about 10% of children and 30% of adults in the United States (US)^4, 5^. NAFLD is the second and most rapidly increasing indication for liver transplantation in the US ^6^.

The spectrum of NAFLD ranges from simple steatosis or nonalcoholic fatty liver (NAFL) with or without mild inflammation, to a more severe phenotype of nonalcoholic steatohepatitis (NASH), which in addition to the presence of steatosis, is characterized by a broad mix of lesions including lobular or portal inflammation, apoptotic hepatocellular injury, and ballooning with or without fibrosis^7^. Different NAFLD phenotypes follow different courses: NAFL without fibrosis rarely causes liver complications while NASH, particularly with fibrosis, carries much higher risk of progression to liver cirrhosis, hepatocellular carcinoma and liver failure, as well as a much higher risk of adverse cardiometabolic outcomes^8-11^.

Similar to the observed rise in NAFLD prevalence in adults^2^, NAFLD prevalence has increased by more than two folds in US adolescents over the past thirty years^12^. As in adults, NAFLD in children is associated with obesity and related cardiac and metabolic comorbidities^13-18^. Despite these similarities, there are unique features and differences in histological phenotypic features between pediatric and adult NAFLD^7, 19-24^. In children, macrosteatosis is usually moderate to severe with panacinar, zone 1 or zone 3 distribution, whereas in adults it is usually mild to moderate in severity with zone 3 or panacinar distribution. While hepatocellular ballooning is the hallmark of adult NASH, it is relatively uncommon in pediatric NASH, particularly before puberty. Children are more likely to have portal inflammation, whereas lobular inflammation predominates in adults. Children are also more likely to have portal-or periportal area-based fibrosis, whereas in adults, early-stage fibrosis is predominantly pericellular or persinusoidal. The molecular basis for these similarities and differences is unclear.

Emerging data suggest that histological characteristics of NAFLD evolve with increasing age in children: following puberty, the severity of steatosis, portal fibrosis and portal inflammation seems to decrease but Mallory-Denk bodies, which are associated with hepatocellular ballooning, start to emerge^23^. Further, after puberty, a shift towards a more adult phenotype may occur^23^, while other children persist with a more periportal pattern of disease. The mechanisms and pathways driving NAFLD progression at different stages in life remain poorly understood.

The variance in histological severity at diagnosis and in patterns of NAFLD progression in children and adults, despite similar clinical risk factors for NAFLD^25-27^, suggests that distinct genetic factors may influence the development, severity, and progression of NAFLD. Hepatic gene expression has been previously studied in adults and to a lesser extent in pediatric populations with NAFLD^25, 26, 28-38^. However, no prior studies directly compared hepatic expression signatures for NAFLD and its sub-phenotypes in well-characterized pediatric and adult patients with liver histology data.

In this study, we aimed to describe molecular mechanisms contributing to similar and discrete NAFLD histopathology at different stages in life by comparing hepatic gene expression profiles for NAFLD and its different phenotypes in well-characterized adolescent and adult patients with obesity. We also sought to determine if transcriptomic signatures of NAFLD phenotypes overlapped between the two age groups and to describe shared pathways between adolescent and adult NASH based on gene ontology analysis.

## Methods

### Study participants

Study subjects included adolescent and adult patients with severe obesity who were undergoing bariatric surgery. Adolescents were recruited through the Teen-LABS study (NCT00474318)^39^, which enrolled consecutive adolescents, aged 19 years and younger, undergoing bariatric surgery (March 2007 to February 2012) at 5 clinical centers in the US: Cincinnati Children’s Hospital Medical Center (Cincinnati, OH), Nationwide Children’s Hospital (Columbus, OH), the University of Pittsburgh Medical Center (Pittsburgh, PA), Texas Children’s Hospital (Houston, TX), and the Children’s Hospital of Alabama (Birmingham, AL), as previously described^37^. Adults were recruited through the bariatric surgery program at the Medical College of Wisconsin between December 2006 and August 2012^26^. The adolescent and adult studies were reviewed and approved by each center’s Institutional Review Board. Written informed consent or assent was obtained from all parents/ guardians and adolescents and directly from adult participants. Standard demographic, clinical and laboratory data were collected on all subjects as previously published^15, 37^.

### Liver histology

A core needle liver biopsy was obtained on all patients during bariatric surgery. Biopsies were scored according to the NASH Clinical Research Network system by an expert pathologist (D.E.K)^40^. The histological phenotypic assignments were: not-NAFLD (which served as controls), NAFL (also referred to as NAFLD-not NASH or simple steatosis), borderline NASH, and definite NASH^40^, based on the aggregate presence and degree of individual histological features of NAFLD.

### Hepatic gene expression analysis

An additional liver biopsy was obtained intraoperatively from each participant and flash frozen for expression studies. Hepatic gene expression was measured using Affymetrix Human Exon 1.0 ST v1 Array (Affymetrix, Santa Clara, CA) in adolescents and Illumina Human-6 Expression Bead Chips (Illumina, Inc. San Diego, CA) in adults. Data were normalized using quantile normalization and baselined to the median of all samples. Differential expression was assessed using an omnibus ANOVA, which compares variability of transcripts within and across diagnostic categories. A fold change requirement of >1.5 for all genes with p<0.05 was considered significant. Pairwise assessments of fold change between diagnoses were constrained to genes that were significant under omnibus testing.

The fold change was assessed in the following comparisons: NAFL vs control, borderline NASH vs control, definite NASH vs control, borderline NASH v NAFL, definite NASH vs borderline NASH, NASH (borderline and definite) vs control, and any NAFLD (NAFL +borderline NASH +definite NASH) vs control. These comparisons were chosen in order to evaluate for transcriptomic independence of NAFL vs controls, borderline NASH vs definite NASH, in addition to NAFLD vs control.

Within each cohort, expression profiles were compared between controls and various NAFLD phenotypes and heatmaps of differentially expressed genes were generated through hierarchical clustering. Differentially expressed genes were compared using Venn diagrams. Principal component analysis (PCA) was used to determine the significant gene set’s differential capacity for identifying NAFLD phenotypes. All analysis was performed in GeneSpring 13.0.

## Results

### Characteristics of study participants

The majority of the adolescent and adult participants were white females (**Table 1**). Adolescents had higher mean body mass index (BMI) (52 ± 9 vs 49 ± 7 kg/m^2^, p=0.01), lower frequency of type 2 diabetes (15% vs 33%, p=0.01) and hypertension (39% vs 58%, p=0.03) but higher frequency of dyslipidemia (73% vs 25%, p<0.0001) compared to adults. Adolescents had higher mean alanine aminotransferase (28±13 vs 22±14 U/L, p=0.03) and aspartate aminotransferase (39±14 vs 21±10 U/L, p<0.0001) levels but lower glucose (97±28 vs 116±45 md/dL, p=0.003), total cholesterol (155±29 vs 175±35 md/dL, p=0.0003), high density lipoprotein (37±8 vs 43±10 mg/dL, p=0.0002), and low density lipoprotein levels (94±27 vs 104±29, md/dL, p=0.02). Although both groups were insulin resistant, there were no significant differences in mean insulin, the homeostatic model assessment for insulin resistance or triglyceride levels between the two groups.

**Table 1.** Characteristics of study participants.

| Cohort | Adolescents<br>n=67 | Adults<br>n=76 | *p* |
| --- | --- | --- | --- |
| Age (years) | 17±1.7 | 43±11.5 | <0.0001 |
| BMI (kg/m<sup>2</sup>) | 52 ± 9.2 | 48.7 ± 7.1 | 0.01 |
| Female n (%) | 52(81.2) | 66(86.8) | 0.1 |
| White n (%) | 49(73.1) | 63(82.8) | 0.1 |
| Type 2 Diabetes n (%) | 10(14.9) | 25(32.8) | 0.018 |
| Dyslipidemia n (%) | 49(73.1) | 19(25) | <0.0001 |
| HTN n (%) | 26(38.8) | 44(57.8) | 0.029 |
| ALT (U/L) | 28±13 | 22±14 | 0.03 |
| AST (U/L) | 39±14 | 21±10 | <0.0001 |
| Glucose (mg/dL) | 97±28 | 116±45 | 0.003 |
| Insulin (μU/mL) | 31±18 | 29±21 | 0.59 |
| HOMA-IR | 7.5±4.6 | 9±8.5 | 0.194 |
| Triglycerides (mg/dL) | 118±62 | 143±100 | 0.08 |
| Total cholesterol (mg/dL) | 155±29 | 175±35 | 0.0003 |
| HDL (mg/dL) | 37±8 | 43±10 | 0.0002 |
| LDL (mg/dL) | 94±27 | 104±29 | 0.02 |

The distribution of histological phenotypes did not differ significantly between adolescents and adults included in this study (p=0.1) (**Figure 1**). NAFLD was present in 49% of adolescents and 61% of adults. The distribution of NAFLD phenotypes between adolescents and adults were NAFL 39% vs 37%, borderline NASH 7% vs 12%, and definite NASH 3% vs 12%, respectively.

**Figure 1.**
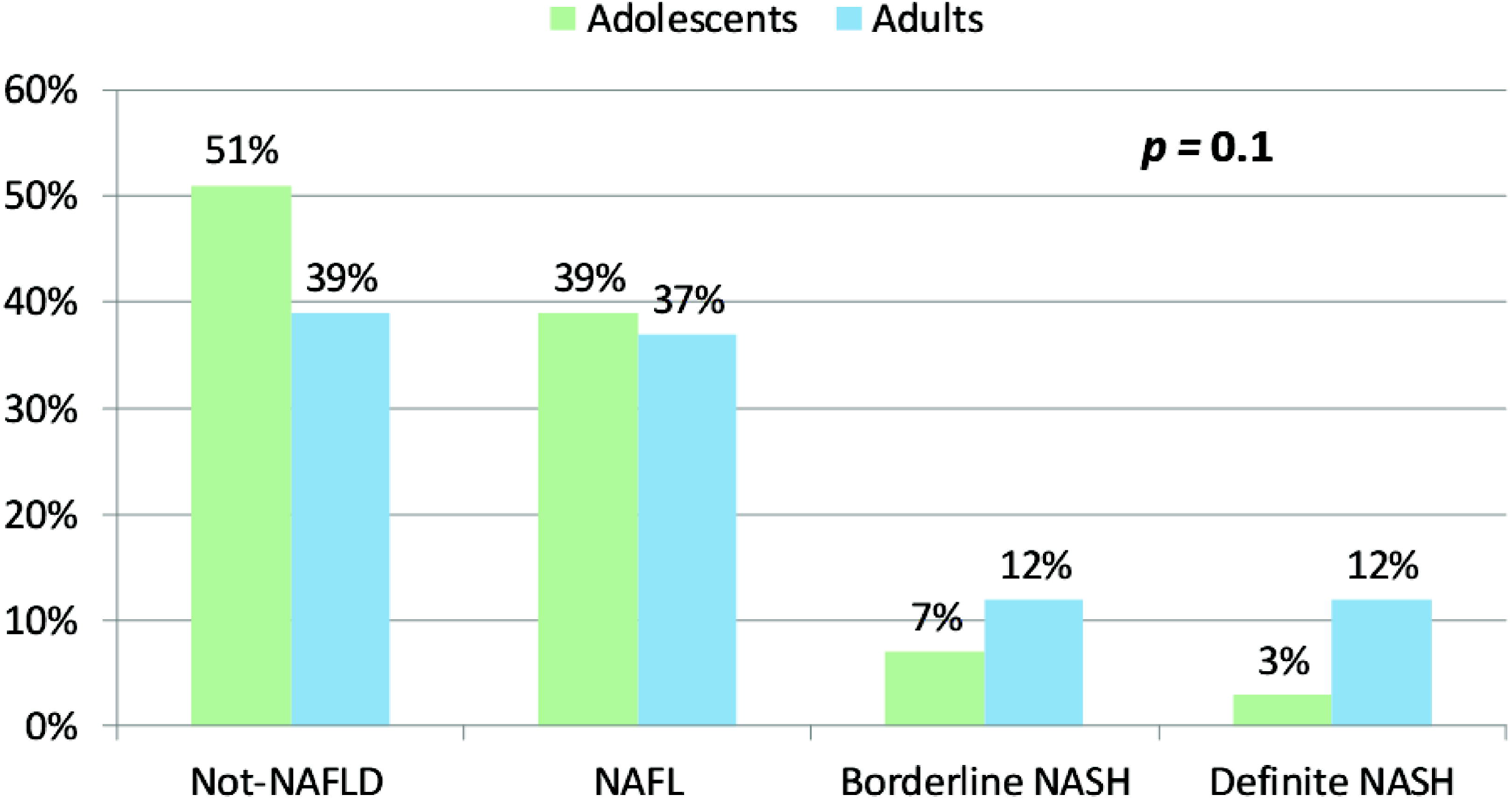
Distribution of histological phenotypes in study participants. Figure 1 footnote: Comparison is between adolescents (n=67) and adults (n=76).

### Hepatic gene expression profiles associated with NAFLD in adolescents and adults

When comparing NAFLD to controls, there were 279 differentially expressed genes in adolescents vs 213 in adults. The heat maps in **Figures 2.A and 3.A** show the clustering of the genes per phenotype in each cohort. The majority of these genes were up-regulated in NAFLD in both cohorts.

**Figure 2.**
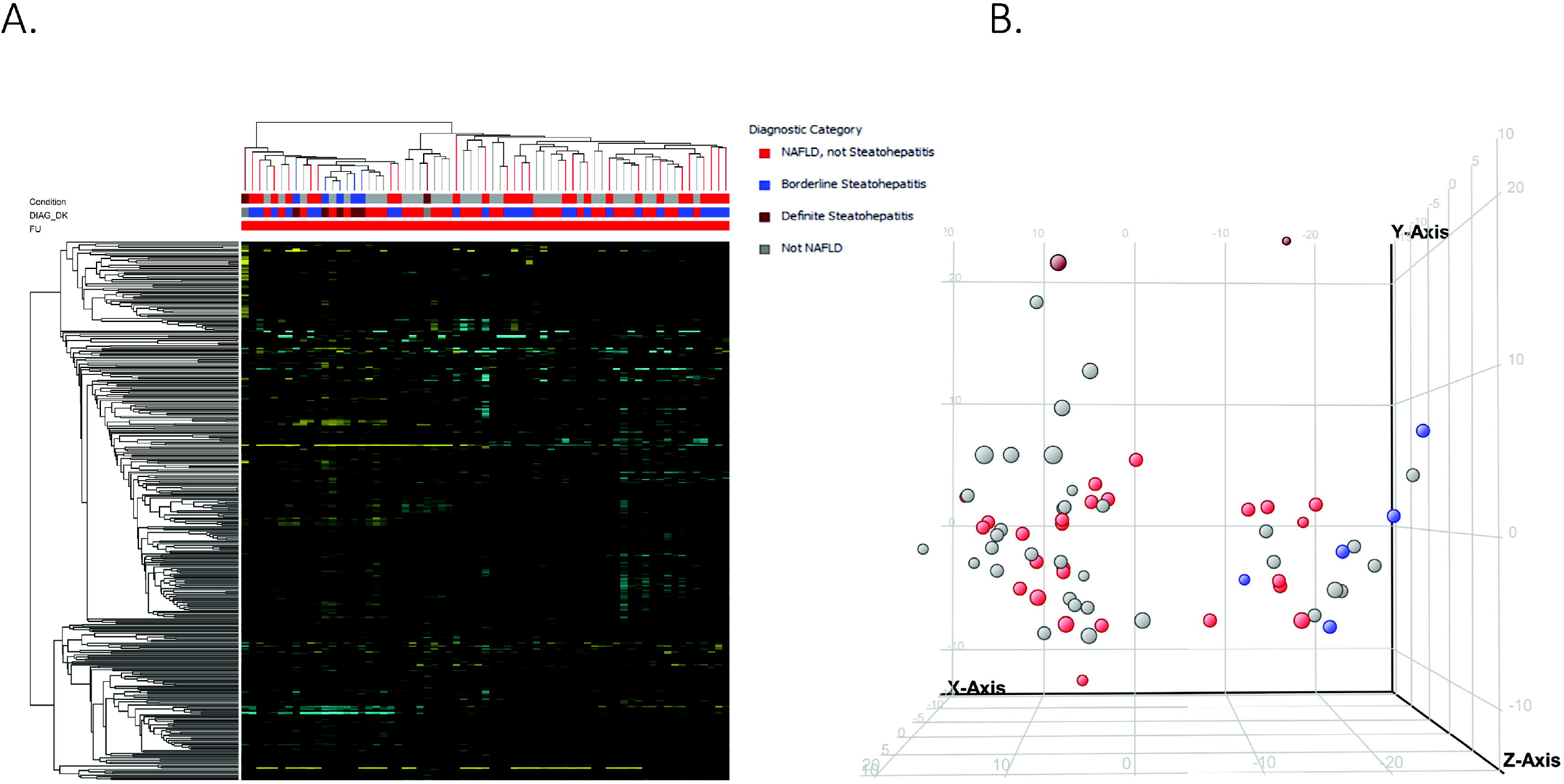
**A**. Heat map and B. Principal component analysis of the 279 differentially expressed genes in adolescents with NAFLD

**Figure 3.**
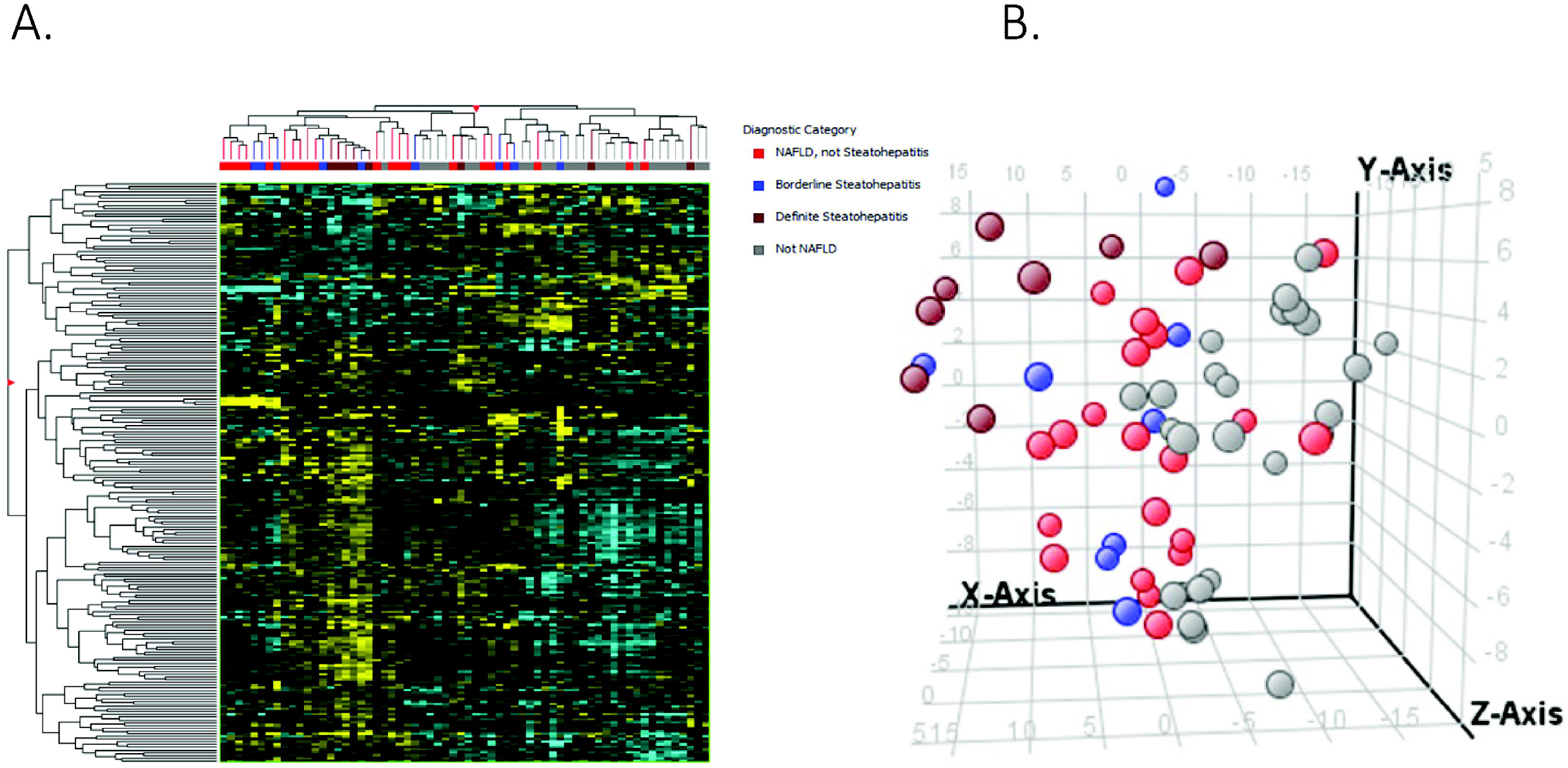
A. Heat map and B. Principal component analysis of the 213 differentially expressed genes in adults with NAFLD

We then performed principal component analysis to assess if the expression changes could differentiate the histological phenotypes. In adolescents, a separation of controls and NAFL from borderline and definite NASH emerged (**Figures 2.B)**. In adults, definite NASH separated from controls, though borderline NASH and NAFL demonstrated heterogeneous distribution and followed a gradient (**Figure 3.B**).

### Portability of differentially expressed NAFLD genes between adolescents and adults

The majority of 279 significant pediatric genes were not portable to the adult population and failed to separate different phenotypes (**Figure 4.A**). Of 279 genes, only 48 (17.2%) were statistically significant in the adult cohort and of those, only 28 (10%) had fold change >1.5. A higher percentage of the 213 significant adult genes were portable to the pediatric population, separating out different phenotypes (**Figure 4.B**). Of 213 genes, 98 (46%) were statistically significant in the pediatric cohort, but only 32 (15%) of those had a fold change >1.5.

**Figure 4.**
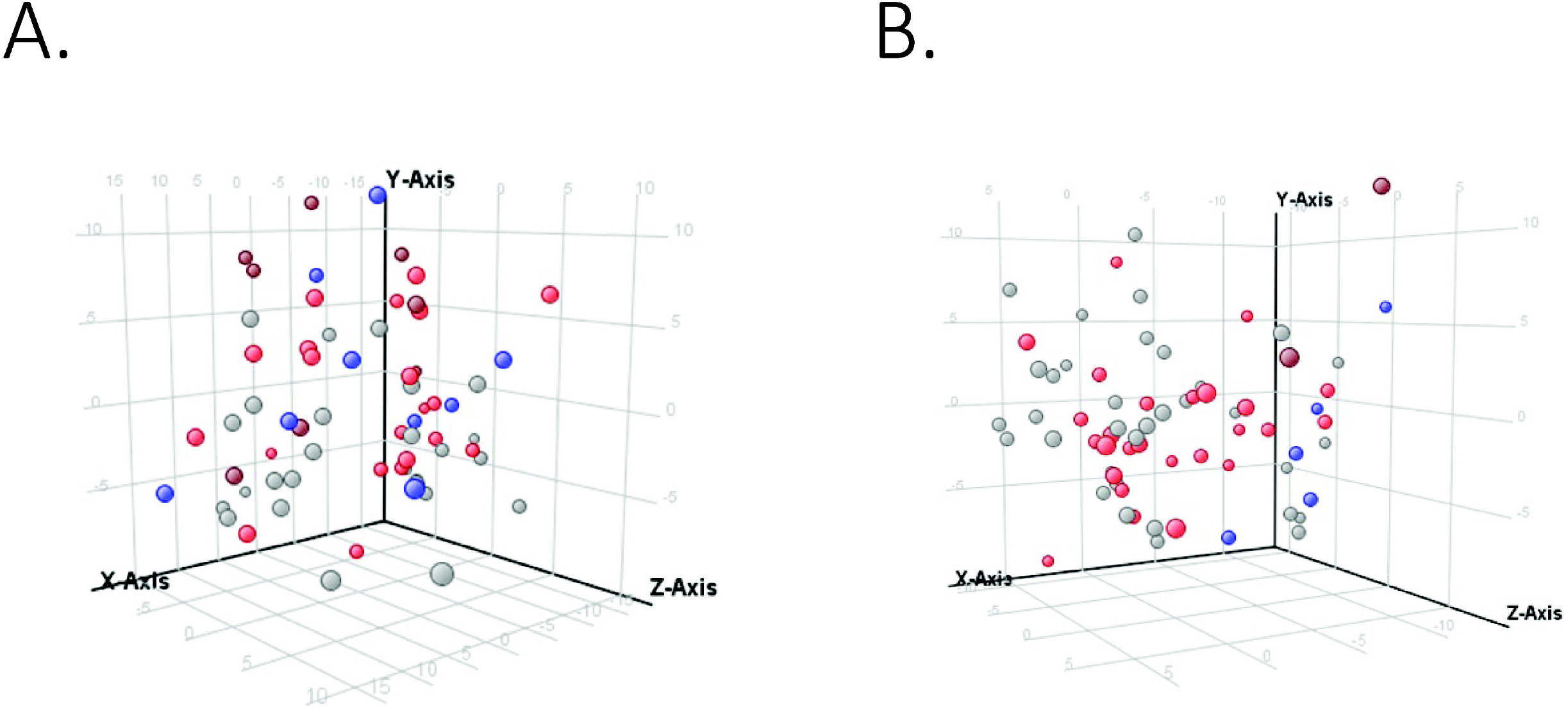
Principal component analysis for portability of differentially expressed NAFLD genes between A) from adolescents to adults and B) from adults to adolescents

### Comparison of expression signatures of different NAFLD phenotypes in adolescents and adults

In adolescents, borderline NASH and definite NASH were transcriptomically similar, although these groups had small numbers number of subjects. In adults, most borderline NASH genes were shared with definite NASH, while the definite NASH gene list had a significant number of genes that were specific to definite NASH (**Figure 5.A**). Pediatric NASH demonstrated more up- and down-regulated genes than adult definite NASH (**Figure 5.B**), albeit this comparison is limited by the small number of patients with pediatric definite NASH. There was very little overlap in differentially expressed genes between pediatric and adult definite NASH: one shared up-regulated gene, complement factor H-related 3 (*CFHR3*) and one shared down-regulated gene, metallothionein 1M (*MT1M*).

**Figure 5.**
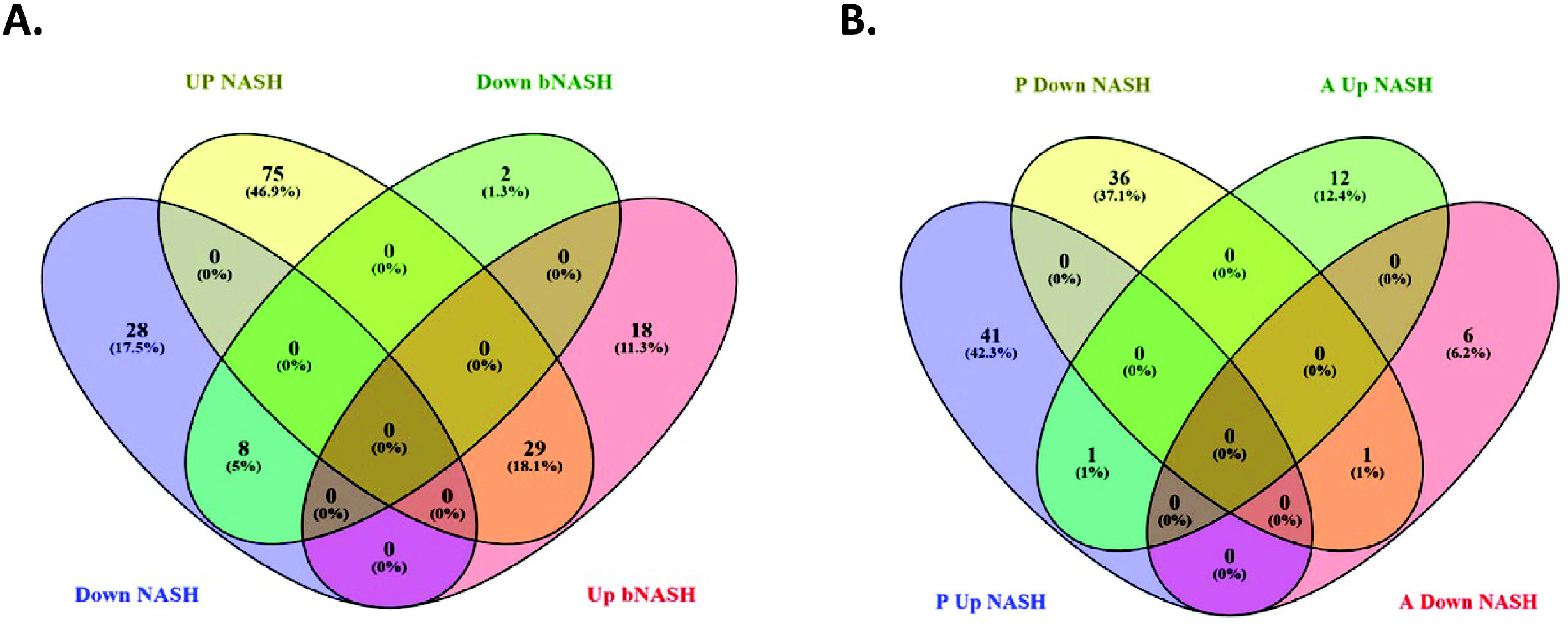
Shared differentially expressed genes between adults with definite and borderline NASH (A) and pediatric and adult NASH (B). Figure 5 footnote: Abbreviations: bNASH: borderline NASH, NASH: definite NASH, P: pediatric, A: adult.

### Gene ontology analysis

When the ontologies of differentially expressed genes were analyzed, borderline NASH in pediatric and adult patients shared consistencies but had notable ontological differences (**Figure 6.A**). For example, gene products upregulated in both included those involved in response to cytokines, increased inflammatory response, PPAR signaling pathway, extracellular matrix organization, response to oxygen containing compounds, and integrin binding, whereas a few shared gene products involved in cysteine-type endopeptidase inhibitor activity, intestinal D-glucose absorption and lysophosphatidic acid binding were downregulated. The glutathione-related functions, biological oxidation, phospholipase A1 inhibitor activity were downregulated only in pediatric borderline NASH, whereas those involved in senescence and autophagy, growth factor binding and wound healing were only upregulated in adult borderline NASH.

**Figure 6.**
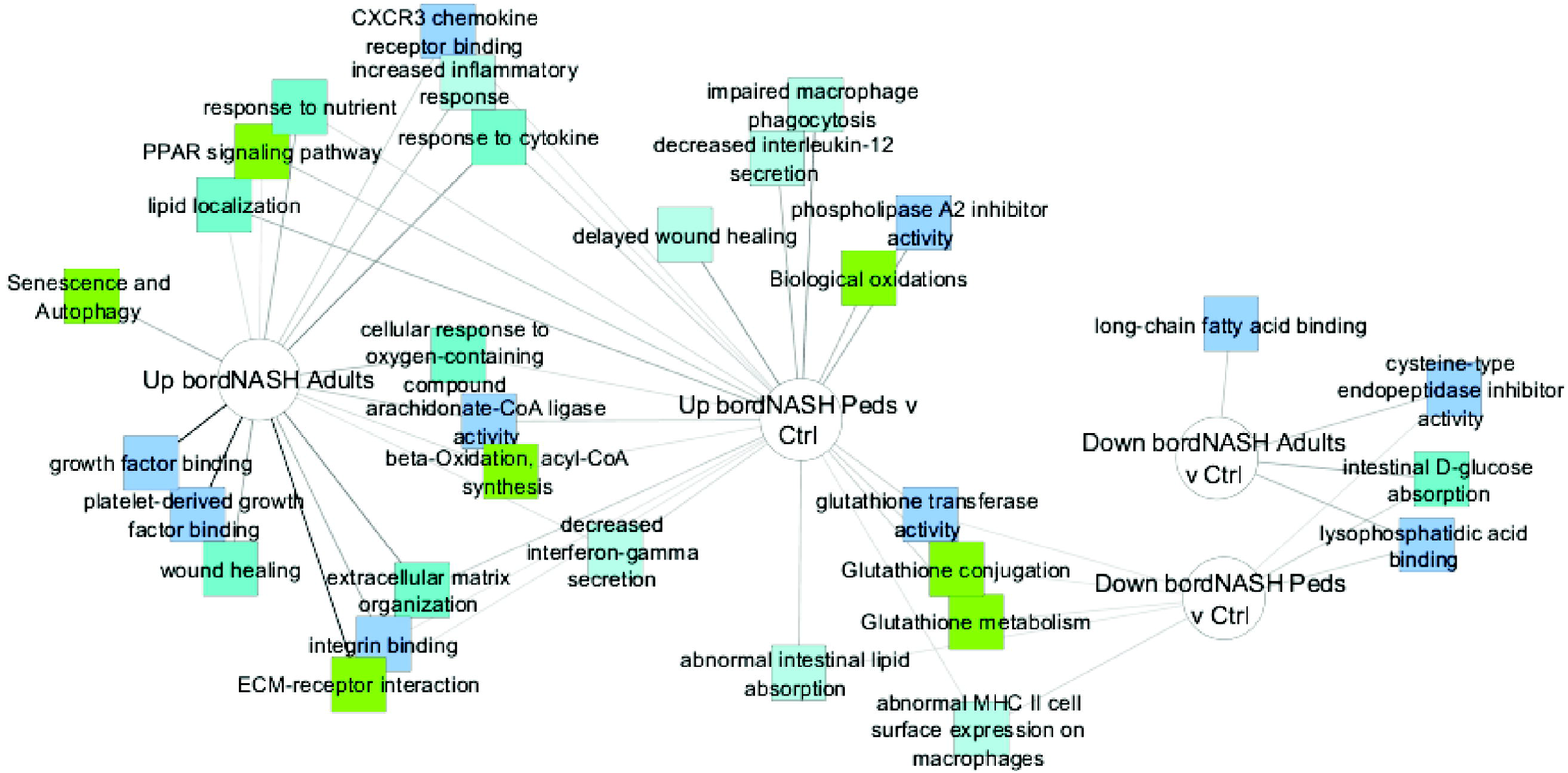
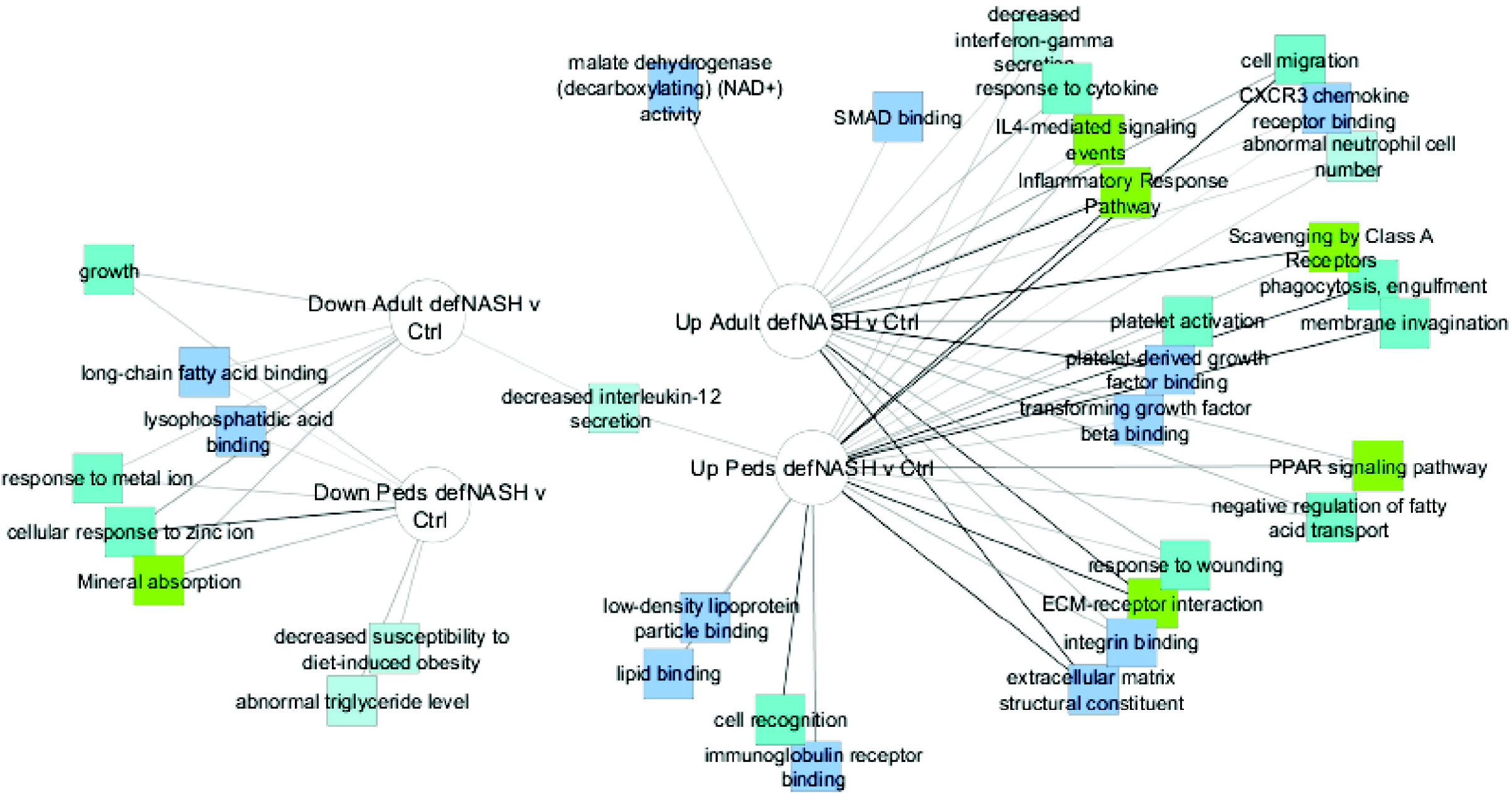
Gene ontology mapping in borderline (A) and definite (B) NASH in pediatric and adult patients Figure 6 footnote: Abbreviations: bNASH: borderline NASH, NASH: definite NASH, P: pediatric, A: adult.

Gene products associated with definite NASH in pediatric and adult patients were consistent ontologically, as shown in **Figure 6.B**. For example, gene products involved in response to cytokine, inflammatory response pathway, cell migration, PPAR signaling pathway, extracellular matrix receptor interaction, integrin binding and extracellular matrix structural constituents were up regulated whereas gene products involved in growth, long-chain fatty acid binding, response to metal ion and mineral absorption were downregulated in both pediatric and adult NASH. Some gene products were uniquely associated with pediatric NASH, which included upregulation of those products involved in low-density lipoprotein particle binding, lipid binding, cell recognition and immunoglobulin binding, and downregulation of those involved in decreased susceptibility to diet-induced obesity and abnormal triglyceride levels.

## Discussion

In this study, we analyzed global hepatic gene expression associated with NAFLD in adolescents and adults who underwent bariatric surgery. NAFLD hepatic expression signatures were different between adolescents and adults. In adolescents, the borderline and definite NASH phenotypes shared similar expression profiles, distinct from the shared genes between pediatric control and NAFL phenotypes. In adults, only definite NASH had a distinctly different expression profile from controls, while NAFL and borderline NASH demonstrated a transcriptomic gradient between controls and definite NASH. Gene ontology analysis showed similarities between pediatric and adult NASH but notable differences suggesting steps leading up to NASH involve different mechanisms in the different age groups.

Hepatic gene expression has been previously studied in adults and to a lesser extent in pediatric populations with NAFLD^25, 26, 28-38^. We previously described global hepatic gene expression in two separate cohorts of adolescent and adult patients with NAFLD who were undergoing bariatric surgery^26, 37^. In adolescents, the fibrogenesis, lipid and carbohydrate metabolism, chemotaxis, cell adhesion, and immune function pathways were upregulated, whereas the glutathione metabolism pathway was significantly downregulated with increased NAFLD severity^37^. In adults, the hepatic fibrosis signaling pathway was upregulated, whereas the endoplasmic reticulum stress and protein ubiquitination pathways were downregulated^26^.

In our comparative analysis, only 17% of the NAFLD genes in adolescents were significant in adults whereas 46% of the adult genes were portable to adolescents. Even smaller percentages of portable genes between age groups had >1.5-fold changes in expression. Gene ontology maps highlighted the shared and different functional categories of NASH-associated differentially expressed genes between adolescents and adults. For example, both age groups shared upregulated genes with functions affecting cell migration, response to cytokines, inflammatory response, platelet activation, PPAR signaling, fibrinogen binding, and extracellular matrix-receptor interaction, and both shared downregulated genes with functions affecting long-chain fatty acid binding, cellular response to zinc and growth. On the other hand, adults had more upregulated genes in unshared functional categories, such as phagocytosis, cell recognition, collagen and lipid bindings and adolescents had more downregulated genes in unshared functional categories involved in glutathione related functions, biological oxidation, phospholipase A1 inhibitor activity, decreased susceptibility to diet-induced obesity and abnormal triglyceride levels were downregulated only in pediatric NASH. Reasons for these differences between adolescents and adults are not clear and the role of other factors influencing NAFLD severity such as sex, hormone levels, exercise and lifestyle patterns at different stages in life need to be further investigated.

Notably, several genome wide association studies (GWAS) in adults and children with NAFLD have identified common single nucleotide polymorphisms (SNPs) associated with increased risk and/or severity of NAFLD (e.g. PNPLA3, TM6SF)^41-46^, and multiple studies have identified unique risk alleles in children, highlighting potential for distinct genetic risk factors associated with earlier onset of disease. A recent GWAS analysis of 208 Hispanic males, age 2-17 year old with well characterized biopsy-confirmed NAFLD identified two novel SNPs, one located within the C8orf17/potassium channel, subfamily K, member 9 (TRAPPC9) gene locus on chromosome 8 that was associated with NAFLD activity score, and one within the ARP5 actin-related protein 5 homolog (ACTR5) gene locus on chromosome 20 that was associated with fibrosis^47^. The role of TRAPPC9 in NAFLD is unknown, while the role of ACTR5 may involve chromatin remodeling.^48^ In another cohort of 118 Italian children with histologically confirmed NAFLD, a SNP in the cannabinoid receptor was associated with severity of inflammation and presence of NASH, though not with steatosis and fibrosis.^49^ A variant in LPIN1 was negatively associated with fibrosis and NAFLD activity score in children, independently of pNLA3 genotype, and clinical risk factors.^50^ LPIN1 is involved in phospholipid and triacylglycerol metabolism.

Our findings likewise suggest potential differences in the pathogenesis and pathways driving severity of NAFLD between pediatric and adult patients despite some shared metabolic and histological similarities ^22^. Therefore, successful therapy of NAFLD may require targeting different pathways in pathogenesis at different stages in life.

This study is the first in-depth analysis of differential gene expression between pediatric and adult patients with NAFLD. The correlation of gene expression was done with well-characterized liver histology as assessed by the same expert liver pathologist who read the pediatric and adult liver biopsies using the same histological assessment system^40^.

This study evaluated select populations of patients with NAFLD. Both adolescents and adults were persons with severe obesity who were undergoing bariatric surgery, and most were white females, which limits the generalization of these findings to other groups with NAFLD not represented in this study. We recognize RNA sequencing is the current and more accurate approach to assess the transcriptome than microarrays. However, microarrays data was already available for the pediatric and adult cohort, together with central histological phenotyping of liver biopsies using the same grading system by the same expert liver pathologist. The availably of these data offered a unique opportunity to try to address the hepatic transcriptome differences between pediatric and adult NAFLD in this study. Expectedly, adolescent had lower frequency of borderline and definite NASH compared to adults, which limits the power of the comparison between these phenotypes. Type 2 diabetes, which has been associated with increased risk of NASH, was also more prevalent among the adult cohort, which may in part account for increased proportion of patients with definite NASH among the adult cohort. However, phenotypic differences in NAFLD were not statistically significant in our study, likely due to smaller sample size.

In summary, this study shows that hepatic gene expression profiles for NAFLD show some similarity and ontological consistency but major transcriptomic differences between pediatric and adult patients with morbid obesity. These data suggest that expression of different genes and pathways contributes to the pathogenesis of NAFLD severity and progression at different stages in life. These findings may have implications for therapeutic drugs target selection in children and adults with NAFLD.

## Abbreviations used

NAFLD: nonalcoholic fatty liver disease
NASH: non-alcoholic steatohepatitis
BMI: Body mass index
AST: Aspartate aminotransferase
ALT: Alanine aminotransferase
HOMA-IR: Homeostatic model assessment for insulin resistance
HDL: high density lipoprotein
LDL: Low density lipoprotein.

## Acknowledgements

Authors thank Fatty Liver Proteomic and Genomic Project Investigators: Harald H.H. Göring (Rio Grande, TX); James Wallace, Mathew Goldblatt, and Richard Komorowski (Milwaukee, WI).

## Notes

**Conflicts of Interests: Dr. Gawrieh:** consulting: TransMedics, Pfizer. Research grant support: Zydus, Galmed, Viking, and Sonic Incytes. **Dr. Chalasani** had paid consulting activities with following companies in last 12 months: Abbvie, Shire, NuSirt, Afimmune, Axovant, Allergan, Madrigal, Coherus, Siemens, and Genentech. He has received research support from Lilly, Galectin, Gilead, Exact Sciences, and Cumberland. Drs. Karn, Kleiner, Olivier declare no conflicts of interest. **Dr. Xanthakos:** consulting: Intercept Pharmaceuticals; research grant support: Target RWE

### Competing Interest Statement

Dr. Gawrieh: consulting: TransMedics, Pfizer. Research grant support: Zydus, Galmed, Viking, and Sonic Incytes. Dr. Chalasani had paid consulting activities with following companies in last 12 months: Abbvie, Shire, NuSirt, Afimmune, Axovant, Allergan, Madrigal, Coherus, Siemens, and Genentech. He has received research support from Lilly, Galectin, Gilead, Exact Sciences, and Cumberland. Drs. Karn, Kleiner, Olivier declare no conflicts of interest. Dr. Xanthakos: consulting: Intercept Pharmaceuticals; research grant support: Target RWE

